# Alteration of actin cytoskeletal organisation in fetal akinesia deformation sequence

**DOI:** 10.1101/2023.06.12.544620

**Authors:** Ramona Jühlen, Valérie Martinelli, Chantal Rencurel, Birthe Fahrenkrog

**Affiliations:** Institute of Molecular Biology and Medicine, Laboratory Biology of the Cell Nucleus, Université Libre de Bruxelles, 6041 Gosselies, Belgium; Institute of Biochemistry and Molecular Cell Biology, Medical School, RWTH Aachen University, 52074 Aachen, Germany; Biozentrum, University of Basel, 4056 Basel, Switzerland

## Abstract

Fetal akinesia deformation sequence (FADS) represents the severest form of congenital myasthenic syndrome (CMS), a diverse group of inherited disorders characterised by impaired neuromuscular transmission. Most CMS originate from defects in the muscle nicotinic acetylcholine receptor, but the underlying molecular pathogenesis is only poorly understood. Here we show that RNAi-mediated silencing of FADS-related proteins rapsyn and NUP88 in foetal fibroblasts alters organisation of the actin cytoskeleton. We show that fibroblasts from two independent FADS individuals have enhanced and shorter actin stress fibre bundles, alongside with an increased number and size of focal adhesions, with an otherwise normal overall connectivity and integrity of the actin-myosin cytoskeleton network. By proximity ligation assays and bimolecular fluorescence complementation, we show that rapsyn and NUP88 localise nearby adhesion plaques and that they interact with the focal adhesion protein paxillin. Based on these findings we propose that a respective deficiency in rapsyn and NUP88 in FADS alters the regulation of actin dynamics at focal adhesions, and thereby may also plausibly dictate myofibril organisation and contraction in skeletal muscle of FADS individuals.

## Introduction

Congenital myasthenic syndromes (CMS) are neuromuscular disorders that primarily affect the neuromuscular junction (NMJ)^1, 2^. The NMJ is a cholinergic synapse by which motor neurons control muscle contraction. Muscle contraction is stimulated by the release of neurotransmitter acetylcholine (ACh) from motor neurons and its binding to the acetylcholine receptor (AChR) at the postjunctional muscle membrane^3^. For stable NMJ formation, AChRs need to cluster and this is regulated by two scaffold proteins: MuSK, a muscle-specific receptor tyrosine kinase, and rapsyn (receptor-associated protein at the synapse). MuSK is activated by binding of agrin, a heparan sulphate proteoglycan, to Lpr4 (low-density lipoprotein receptor-related protein 4), and by binding of DOK7 (downstream of tyrosine kinase 7). Full activation of MuSK results in activation of rapsyn, an increase in rapsyn’s concentration at the NMJ, and in consequence in clustering of AChRs^2^. Biallelic mutations in the genes coding for MuSK, rapsyn, and DOK7 cause CMS, in particular fetal akinesia deformation sequence (FADS), the severest form of CMS^4–9^. FADS is an aetiologically heterogenous condition which is distinguished by the inability of affected foetuses to initiate movement *in utero* ^10, 11^. For those affected, the lack of foetal movement is responsible for distinct developmental defects, including joint contractures and lung hypoplasia. FADS foetuses are predominantly premature or stillborn, and live-births have a high mortality rate due to respiratory failure.

Beyond the MuSK-DOK7-rapsyn signalling axis, biallelic mutations in the gene coding for nucleoporin 88 (NUP88) were recently identified as cause for FADS^12^. Loss of NUP88 function resulted in defects in AChR maturation in zebrafish and reduced expression of rapsyn^12^. In accordance with the notion that signalling downstream of MuSK and rapsyn involves interactions with the cytoskeleton, we recently reported that depletion of rapsyn and NUP88 perturbed the microtubule network. This perturbation led to defective primary cilium formation and impaired ciliogenesis^13^. The defects in ciliogenesis may account for the pleiotropic developmental defect seen in FADS, but are likely not the basis for defective AChR clustering. Clustering of AChRs, however, is orchestrated by the actin cytoskeleton and by actin dynamics^14–19^. Here, we provide evidence that fibroblasts from FADS individuals are characterised by multiple alterations in actin cytoskeleton organisation, including enhanced actin-myosin bundling, enlarged size and number of focal adhesions, and elevated levels of RhoA. We show that the FADS-related proteins rapsyn and NUP88 interact with the focal adhesion protein paxillin at adhesions plaques and that FADS-related mutations in rapsyn and NUP88 disturb this interaction. In light of our results, we propose that dysregulation of the actin cytoskeleton due to the loss of regulatory function of FADS-related proteins at focal adhesions contributes to FADS pathology.

## Results

### Rapsyn and NUP88 deficiency coincide with altered actin stress fibre appearance

We have recently shown that the formation of primary cilia was compromised in fibroblasts derived from FADS individuals. Likewise, depletion of the FADS-related proteins rapsyn and NUP88 interfered with primary cilia formation^13^. A cross-talk between ciliogenesis and actin cytoskeleton dynamics is well described^20–23^, which prompted us to investigate the effect of FADS-like changes on the actin cytoskeleton. To do so, we first depleted rapsyn or NUP88 from normal human foetal MRC5 fibroblasts by RNA interference. Forty-eight hours after siRNA transfection, the actin cytoskeleton was visualised by fluorophore-labelled phalloidin (phalloidin-Alexa 488) and examined by confocal microscopy. We found that actin stress fibres (SFs) appeared reduced upon the respective depletion of rapsyn and NUP88 from MRC5 cells (Fig. 1a). This effect was importantly also evident in primary skin fibroblasts from two FADS individuals, referred to as FADS 1 and FADS 2 (Fig. 1b). FADS 1 fibroblasts were derived from a foetus carrying an unknown mutation (Coriell Institute; ID: GM11328) and FADS 2 fibroblasts harbour the missense mutation p.E162K in *RAPSN*^4, 13^. Consistent with alterations in actin SFs, FADS 1 and FADS 2 fibroblasts exhibited a more elevated and polygonal shaped morphology, as compared to the flat, elongated and spindle-like shaped MRC5 control fibroblasts (Fig. 1c).

**Figure 1.**
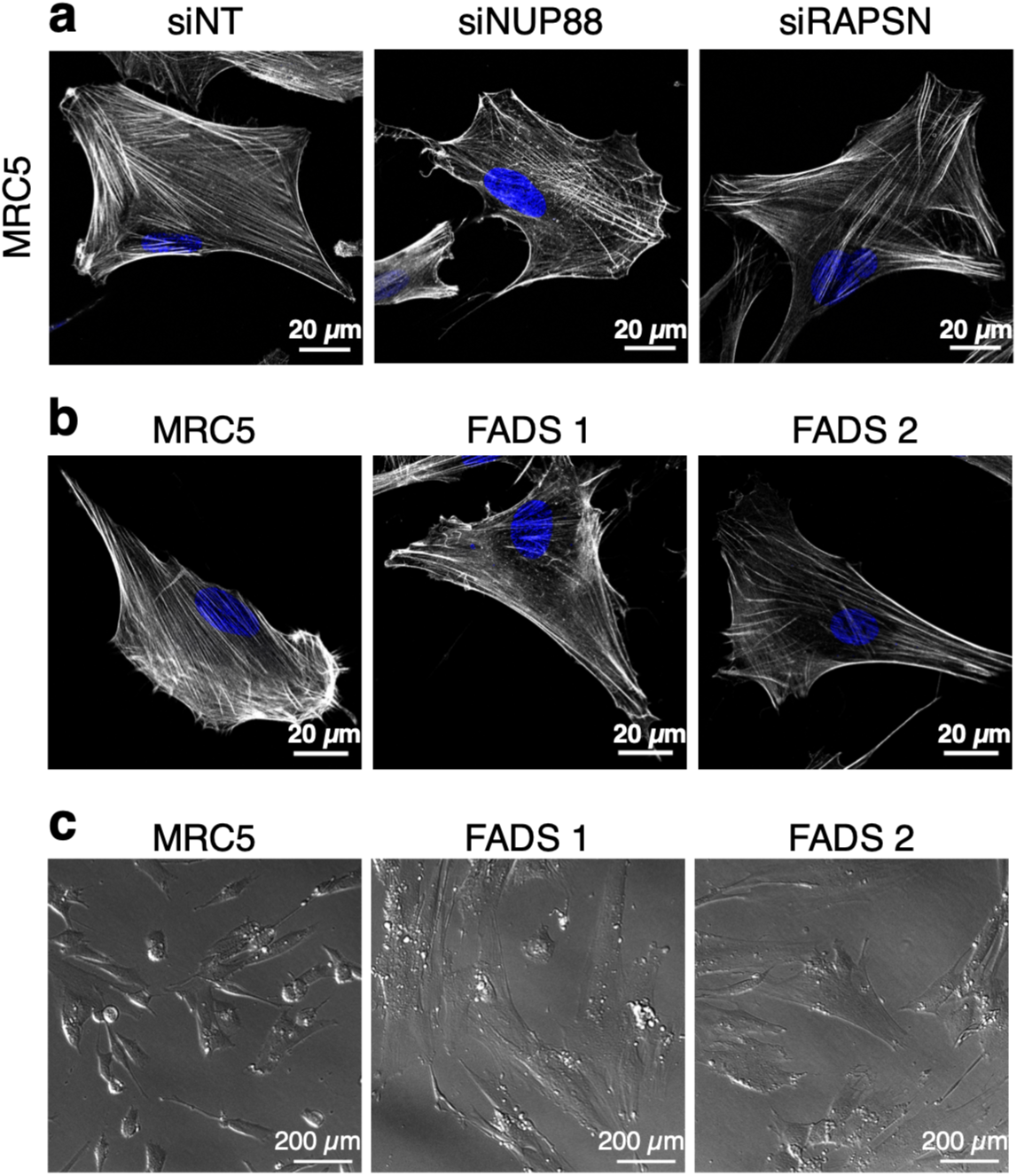
Actin stress fibre formation is perturbed in FADS. (**a**) Actin stress fibres appeared reduced in MRC5 cells NUP88-(siNUP88) and rapsyn (siRAPSN)-depleted as compared to cells treated with non-targeting siRNAs (siNT) and (**b**) in fibroblasts derived from two FADS individuals. Cells were stained with phalloidin to visualise F-actin (greyscale) and DAPI to visualise DNA (blue). Shown are representative confocal images. (**c**) Representative transmitted light images of MRC5 and FADS fibroblasts grown on glass coverslips.

### Altered cytoskeletal organisation and increased actin-myosin bundling in FADS fibroblasts

To analyse the differences in the actin cytoskeleton in a more quantitative way, we next took advantage of crossbow-shaped surface micropatterns. Such adhesive surface micropatterns induce a controlled polarity of the cytoskeleton due to which cell shape and morphology become normalised^24^. The actin cytoskeleton is very dynamic and can organise itself distinctively (Fig. 2a): branched actin characterises lamellipodia at the leading edge of the cell, parallel non-contractile actin bundles are found in dorsal SFs, whereas anti-parallel contractile actin-myosin bundles form transverse actin arcs and ventral SFs. Dorsal and ventral SFs connect to focal adhesions (FAs), which attach the extracellular matrix to the cytoskeleton via integrins, and are mainly anchored to the actin cytoskeleton by different mechano-sensing linkers, such as vinculin^25–27^. When grown on crossbow-shaped micropatterns, cells typically exhibit a stereotypic organisation of the actin cytoskeleton: lamellipodia appear at the curved leading edge of the cell, dorsal SFs are perpendicular to the leading edge and attach to transverse arcs that line the leading edge. Ventral SFs flank the centre of the crossbow at the trailing edge (Fig. 2a)^28, 29^. To particularise the organisation of the actin cytoskeleton in FADS fibroblasts, we first visualised actin SFs by phalloidin and contractile actin-myosin bundles using an antibody to non-muscle myosin II A (NMIIA). Their respective distribution was analysed automated using an ImageJ plug-in and results were illustrated by a colour-coded frequency map denoting pixel intensities (Fig. 2b, Supplementary Fig. S1 online). We found that contractile transverse arcs and ventral SFs were augmented, whereas non-contractile dorsal SFs were reduced in FADS fibroblasts (Fig. 2a,b). Transverse arcs and ventral SFs are characterised by actin-myosin bundling promoted by α-actinin, which cross-links F-actin and is especially abundant on SFs (Crawford et al., 1992). Consistently, the distribution of α-actinin resembled the phalloidin and NMIIA staining patterns (Fig. 2b). The increase in actin-myosin bundles in FADS cells was not due to changes in the frequency of NMIIA peaks (Supplementary Fig. S2a,b online). Moreover, the overall orientation of actin filaments within the cell (Fig. 2c, phalloidin), as well as the number of actin branches and junctions (Supplementary Fig. S2c,d online), remained unaltered in FADS cells, indicating that the overall connectivity and integrity of the actin network did not change.

**Figure 2.**
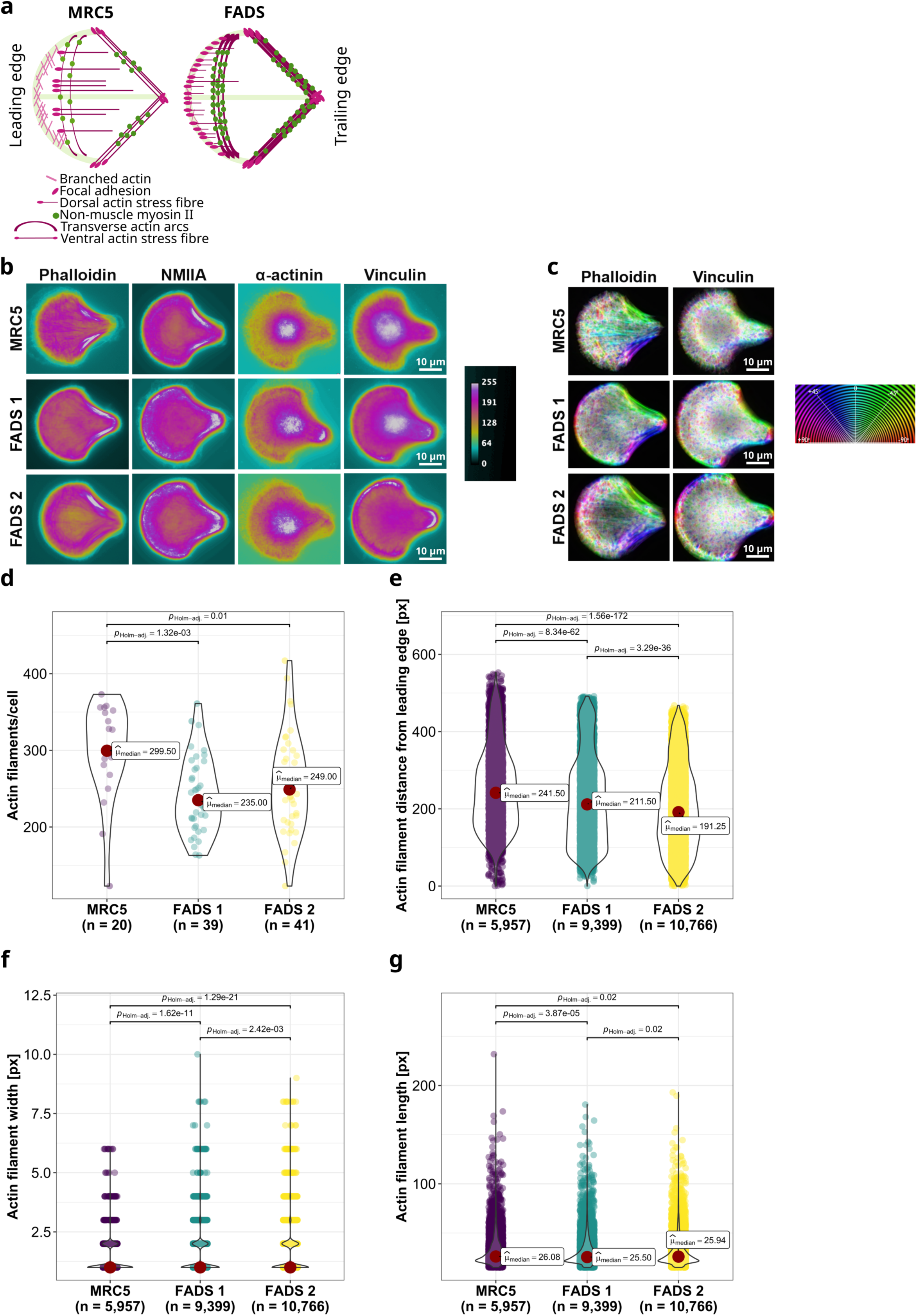
FADS fibroblasts show localised alterations in actin cytoskeletal organisation and enhanced actin stress fibre bundling. (**a**) Schematic summary depicting typical actin organisation in MRC5 control and in FADS fibroblasts. FADS fibroblasts have augmented contractile transverse arcs and ventral stress fibres. Dorsal stress fibres and focal adhesions are shifted to the leading edge of FADS fibroblasts. (**b**) Colour-coded frequency maps highlight local rearrangements of the actin cytoskeleton in FADS fibroblasts as compared to MRC5 control cells. Cells were grown on crossbow-shaped micropattern and analysed by immunofluorescence microscopy. Images were processed as illustrated in the analysis workflow shown in Supplementary Fig. S1 online. Phalloidin was used to visualise F-actin, non-muscular myosin II A (NMIIA) and actinin were used to delineate contractile actin-myosin bundles, and vinculin was used to display mature focal adhesions. The colour-coded map indicates pixel intensities from 0 to 255. (**c**) Orientation of F-actin (phalloidin) and mature focal adhesions (vinculin) in MRC5 and FADS fibroblasts. The respective distribution of F-actin (phalloidin) and vinculin was sampled from images as shown in (b). The circular colour-coded map shows their respective orientation. (**d-g**) Quantitative analysis of actin stress fibres using the open-source JRE analysis tool FilamentSensor. Shown are violin plots summarising the analysis of: (**d**) actin filament number per cell, (**e**) the distance of actin filaments from the cells’ leading edge, as well as (**f**) the width and (**g**) the length of actin filaments. Dunn’s non-parametric test was used to calculate pairwise statistics for Kruskal-type ranked data (*p*_Holm-adj._). n denotes the number of observations and µ the median (red dot). px, pixel.

To further quantify the changes in actin organisation in FADS cells, we employed the FilamentSensor tool, which allows to define location, orientation, length, and width of single filaments^30^. We found that the overall number of actin filaments was reduced in FADS fibroblasts (Fig. 2d), that actin filaments were shifted towards the curved leading edge of the cell (Fig. 2e), and that actin filaments were generally thicker and shorter in FADS cells (Fig. 2f,g). Together this suggests that an increased actin-myosin bundling results in a gain of thicker actin filaments and in turn a reduction in the overall number of individual actin filaments in FADS fibroblasts.

### Distinct rearrangement of focal adhesions in FADS fibroblasts

Given the increase in ventral SFs in FADS fibroblasts, we next examined vinculin, which connects ventral stress fibres to distant FAs^25, 27^. Similar to ventral SFs, FAs were likewise increased in FADS cells, both at the leading and the trailing edge of the cell and their orientation within the cell resembled those of the SFs (Fig. 2a-c). Interestingly, while the width of FAs was increased in both FADS fibroblast lines (Fig. 3a), their overall number was only augmented in FADS 2 cells (Fig. 3b). Furthermore, mapping the respective distribution of thick actin filaments and thick FAs in FADS cells revealed that thick actin filaments were enriched in transverse actin arcs and ventral SFs (Figs. 2a, 3c), while thick FAs were shifted towards the leading edge of the cell as part of dorsal SFs and to the trailing edge as part of ventral SFs (Figs. 2a, 3c).

**Figure 3.**
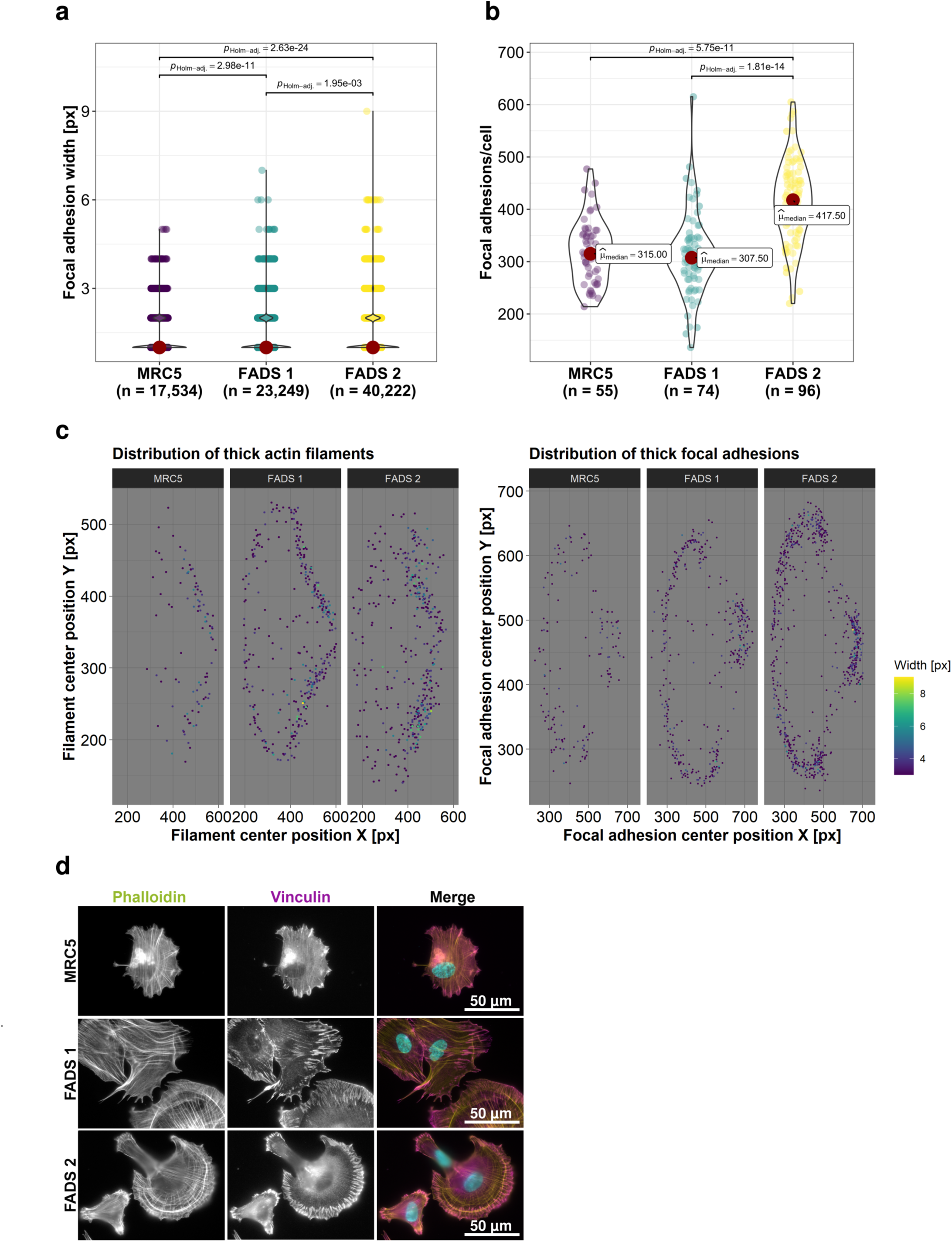
FADS fibroblasts have altered focal adhesion and spreading properties. (**a-b**) Quantitative analysis of focal adhesions in MRC5 and FADS fibroblast grown on crossbow-shaped micropatterns using the FilamentSensor tool. Shown are violin plots summarising the analysis of: (**a**) the width of focal adhesions and (**b**) their number per cell. Dunn’s non-parametric test was used to calculate pairwise statistics for Kruskal-type ranked data (*p*_Holm-adj._). n denotes the number of observations and µ the median (red dot). In (a) the median descriptive were omitted because here all values are 1. (**c**) Colour map representations of the localisation of thick actin filaments (left) and thick focal adhesions (right) in MRC5 and FADS fibroblasts. The x-axis indicates the X-coordinate of the filament or adhesion centre and the y-axis the Y-coordinate of the filament or adhesion centre within the crossbow-shaped micropattern. The colour map resembles the filament or adhesion width in pixel (px). Thick actin filaments and thick focal adhesions are mainly found within contractile transverse arcs and ventral stress fibres in FADS fibroblasts (see also Fig. 2a). (**d**) Representative epifluorescence images of fibroblasts imaged 3 hours after seeding on crossbow-shaped micropattern. FADS fibroblasts show enhanced spreading immediately after seeding and developed characteristic membrane protrusions. Phalloidin (green) was used to visualise F-actin, vinculin (magenta) to depict mature focal adhesions, and DAPI (blue) to stain DNA.

FAs promote plasma membrane protrusions at the leading edge of migrating cells by inhibiting the retrograde actin flow and by converting actin-myosin pulling into traction^31^. The described rearrangements of FAs in FADS cells therefore suggest that cell adhesion and/or motility might be impaired. Consistently, FADS cells adhered faster and developed pronounced adhesive FAs and contractile SFs as compared to MRC5 cells (Fig. 3d), when analysed 3 hours after seeding. Together these findings suggest that FADS fibroblasts exhibit enhanced forces through FAs and actin-myosin bundles to drive forward movement by membrane protrusions.

### Localised myosin contraction in FADS cells is induced by enhanced Rho activation

We have noticed that FADS fibroblasts, when grown on surface micropatterns, developed pronounced plasma membrane blebs at the leading and trailing edge. The bleb cortex stained positive for F-actin (phalloidin), NMIIA, and vinculin (Fig. 4a, arrowheads). The cortex and the interior of membrane blebs were also positive for phosphorylated regulatory light chain of myosin (MLC-P; Fig. 4b, arrowheads), as well as the outer edges of the cell. MLC-P is prerequisite for bundling of contractile SFs and given the enhanced actin-myosin bundling in FADS cells (Fig. 2f,g), this suggests that membrane blebbing is a result of an increased intracellular pressure due to altered actin-myosin contraction^32^.

**Figure 4.**
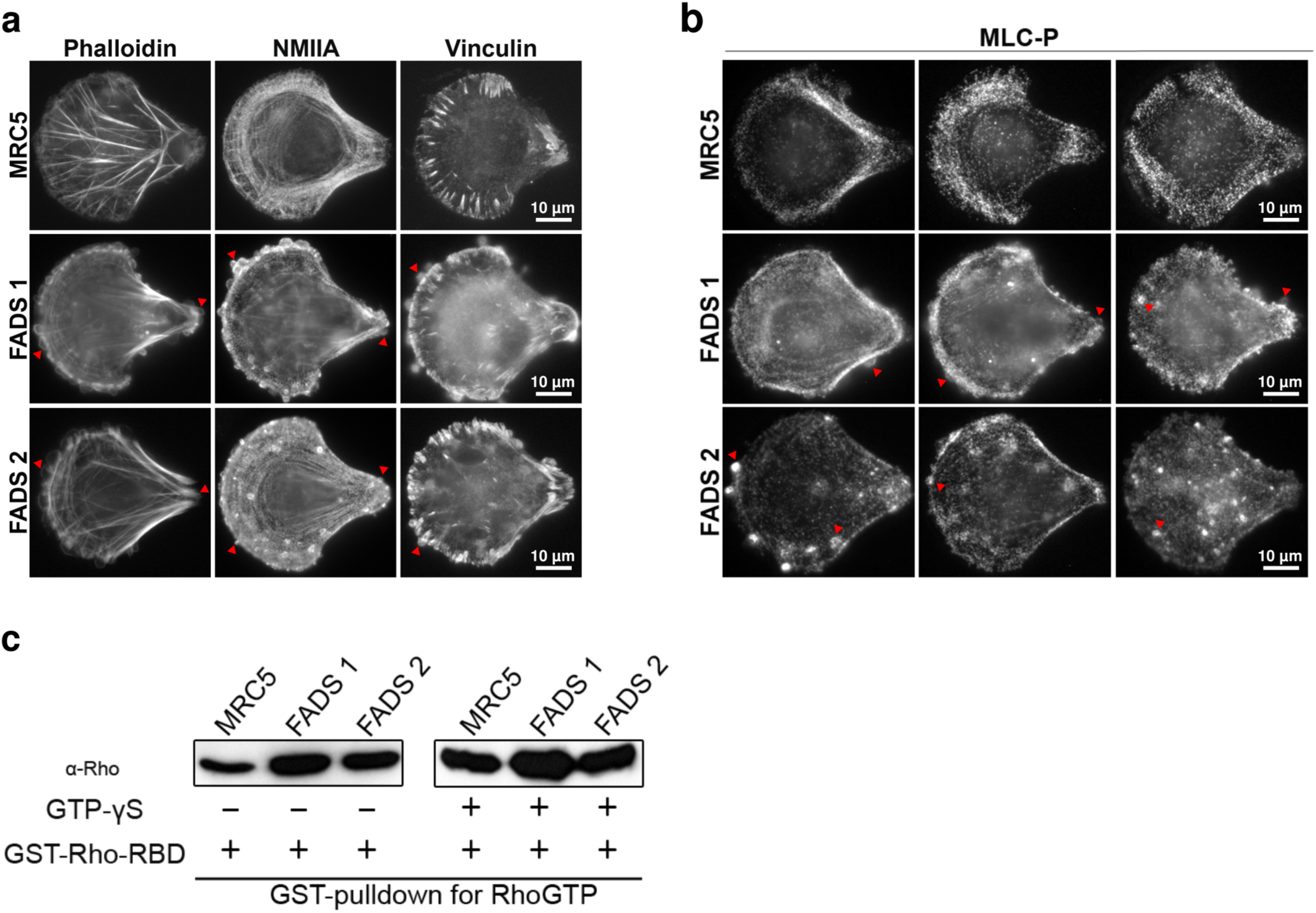
FADS fibroblasts develop plasma membrane blebs and have increased Rho activity. Representative epifluorescence images of MRC5 and FADS fibroblasts grown on crossbow-shaped micropattern, (**a**) stained for F-actin (Phalloidin), non-muscular myosin II A (NMIIA) and vinculin, as well as (**b**) phosphorylated regulatory light chain of myosin (MLC-P). Membrane blebs at the leading and trailing edge in FADS fibroblast are marked with red arrow heads. Bleb cortex are positive for F-actin, NMIIA and vinculin, whereas bleb interiors are positive for MLC-P. MLC-P staining is enhanced at the edges of FADS fibroblasts. (**c**) GST-pulldown for active Rho revealed that FADS fibroblasts have enhanced RhoGTP levels as compared to MRC5 cells. GTP-γS is a non-hydrolysable GTP-analogue. GST-Rho-RBD, GST-Rhotekin-Rho binding domain.

Phosphorylation of MLC is mediated by distinct serine/threonine kinases which in turn are regulated by Rho GTPases. In this context, RhoA regulates actin-myosin bundling by activating the ROCK kinase and the formin mDia1^33, 34^. By using a GST-pulldown for RhoGTP, we found that the levels of active Rho were increased in FADS cells as compared to control cells (Fig. 4c, left blot). When adding the non-hydrolysable GTP-analogue GTP-γS to the assay to activate Rho, the levels of RhoGTP increased in the control cells and were comparable to those in FADS cells (Fig. 4c, right blot). These data indicate that FADS fibroblasts exhibit an increased level of active Rho, which likely directly enhances actin-myosin bundling and contraction. By a sliding mechanism, increased actin-myosin bundling and contraction provokes wider and shorter actin filaments, exactly as described above (Fig. 2f,g).

### FADS-related proteins rapsyn and NUP88 are nearby adhesion plaques

To further explore focal adhesions in context of FADS, we next performed *in situ* proximity ligation assays (PLA) in MRC5 and FADS fibroblasts (see Materials and Methods)^35^. We analysed the proximity between rapsyn and NUP88, and vinculin. In MRC5 cells, about 10 foci per cell were detected for vinculin:rapsyn (Fig. 5a,b) and about 15 per cell for vinculin:NUP88 (Fig. 5c,d). In FADS 1 cell,s the number of foci increased significantly for both vinculin:rapsyn (about 50 foci per cell) as well as vinculin:NUP88 (about 110 foci per cell) and likewise in FADS 2 cells for vinculin:NUP88 (about 50 foci per cell). The number of PLA foci for vinculin:rapsyn in contrast was reduced in FADS 2 fibroblasts (about 4 foci per cell), likely due to the point mutation in rapsyn in these cells. PLA results were validated by negative controls (Supplementary Fig. S3 online). Together our data suggest that the association of rapsyn and NUP88 with vinculin is perturbed in FADS cells.

**Figure 5.**
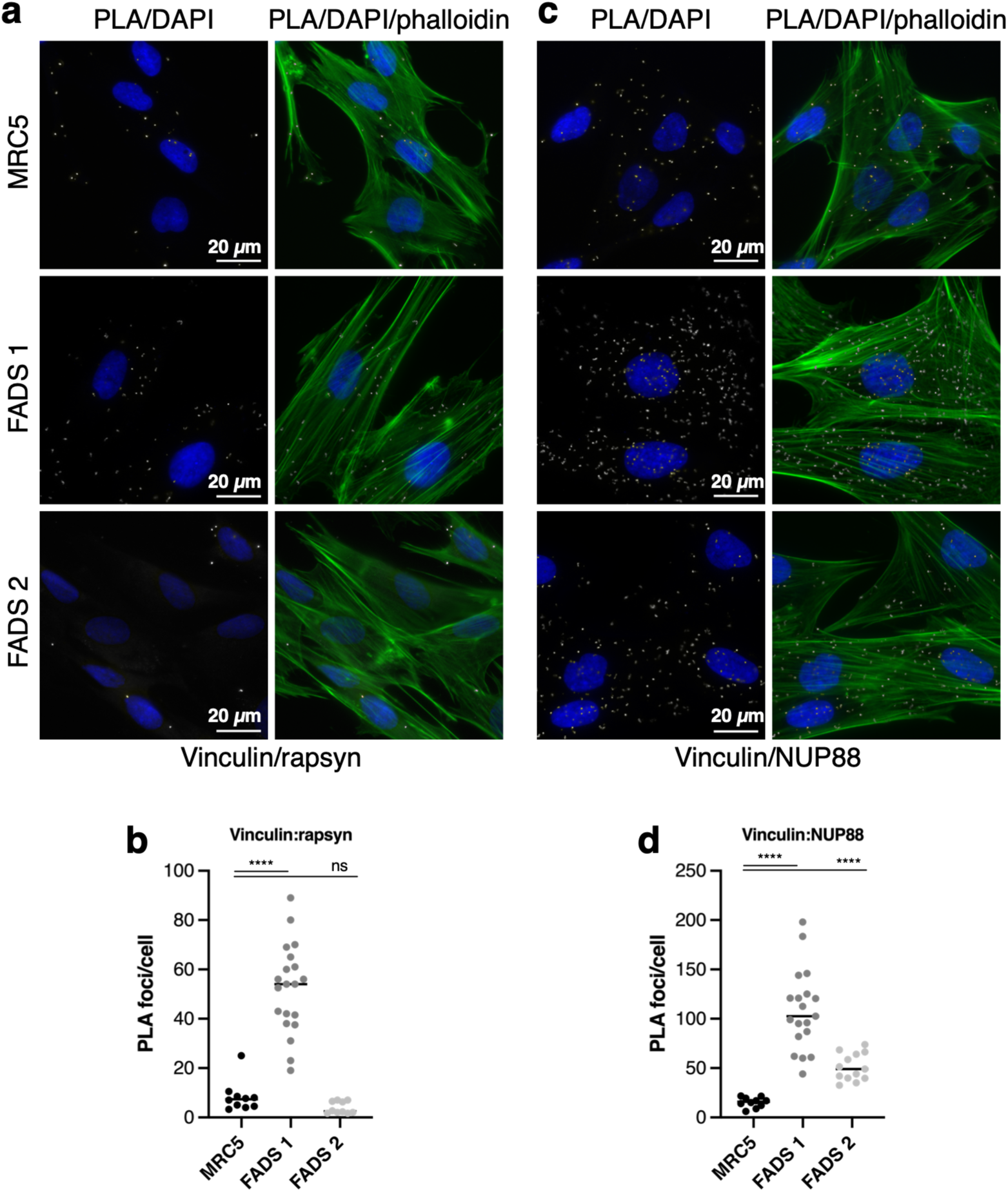
The association of rapsyn and NUP88 with vinculin is altered in FADS fibroblasts. (**a**) Close proximity (about 40 nm) between vinculin and rapsyn was visualised by PLA foci (grey) and (**b**) quantified. Total number of analysed cells: MRC5, n=21; FADS 1, n=25; FADS2, n=36. (**c**) PLA association between vinculin and NUP88 and (**d**) quantification of PLA foci per cell. Total number of analysed cells: MRC5, n=41; FADS 1, n=36; FADS 2, n=60. Phalloidin was used to visualise F-actin (green) and DAPI (blue) for DNA. Shown are representative immunofluorescence images from at least three independent experiments. Black horizontal line represents the median. *\*\*\*\*p* < 0.0001; ns, *p* not significant. Two-Way Anova test was used to calculate statistics.

For further assessment of rapsyn and NUP88 association with focal adhesions, we exploited the bimolecular fluorescence complementation (BiFC) system in HeLa cells. BiFC is based on protein interactions bringing together ectopically expressed N-terminal (V1-(or VN-)) and C-terminal (V2-(or VC-)) fragments of the Venus protein to reconstitute fluorescence, thus allowing direct visualisation of protein interactions in their normal cellular environment^36^. Rapsyn is known to bind to several actin-binding proteins^37–39^, hence we tested the system demonstrating that expression of rapsyn-V1 and actin-V2 in HeLa cells resulted in strong fluorescence in the cytoplasm (Fig. 6a; Supplementary Table S1). In contrast, NUP88 paired with actin did not produce a fluorescence signal (Supplementary Fig. S4a online). The fluorescence signal arising from the rapsyn:actin pair was used as reference signal for adjustment of the laser intensity on the confocal microscope to allow quantitative analysis of the assays. The FADS-related E162K mutant of rapsyn paired with actin resulted in cytoplasmic fluorescence similar to the wild-type rapsyn:actin (Fig. 6a). Important for the interaction of actin with actin-binding proteins is a hydrophobic cleft between subunit 1 and 3 of actin^40^, and mutations affecting this cleft, i.e. actinΔ143-146 (Fig. 6b) and actinΔ346-349 (Supplementary Fig. S4a online) led to a significant decrease in the cytoplasmic fluorescence of the rapsyn:actin pair (Fig. 6b). Mutations that affect actin polymerisation^41^ (actin G215D; Fig. 6b) or mutations in residues surrounding the hydrophobic cleft (actin E167A and actin S350A; Supplementary Fig. S4a online) on the other hand did not impair reconstitution of the Venus fluorescence. Thus, we conclude that rapsyn interacts with actin and that the interaction does not depend on the polymerisation status of actin.

**Figure 6.**
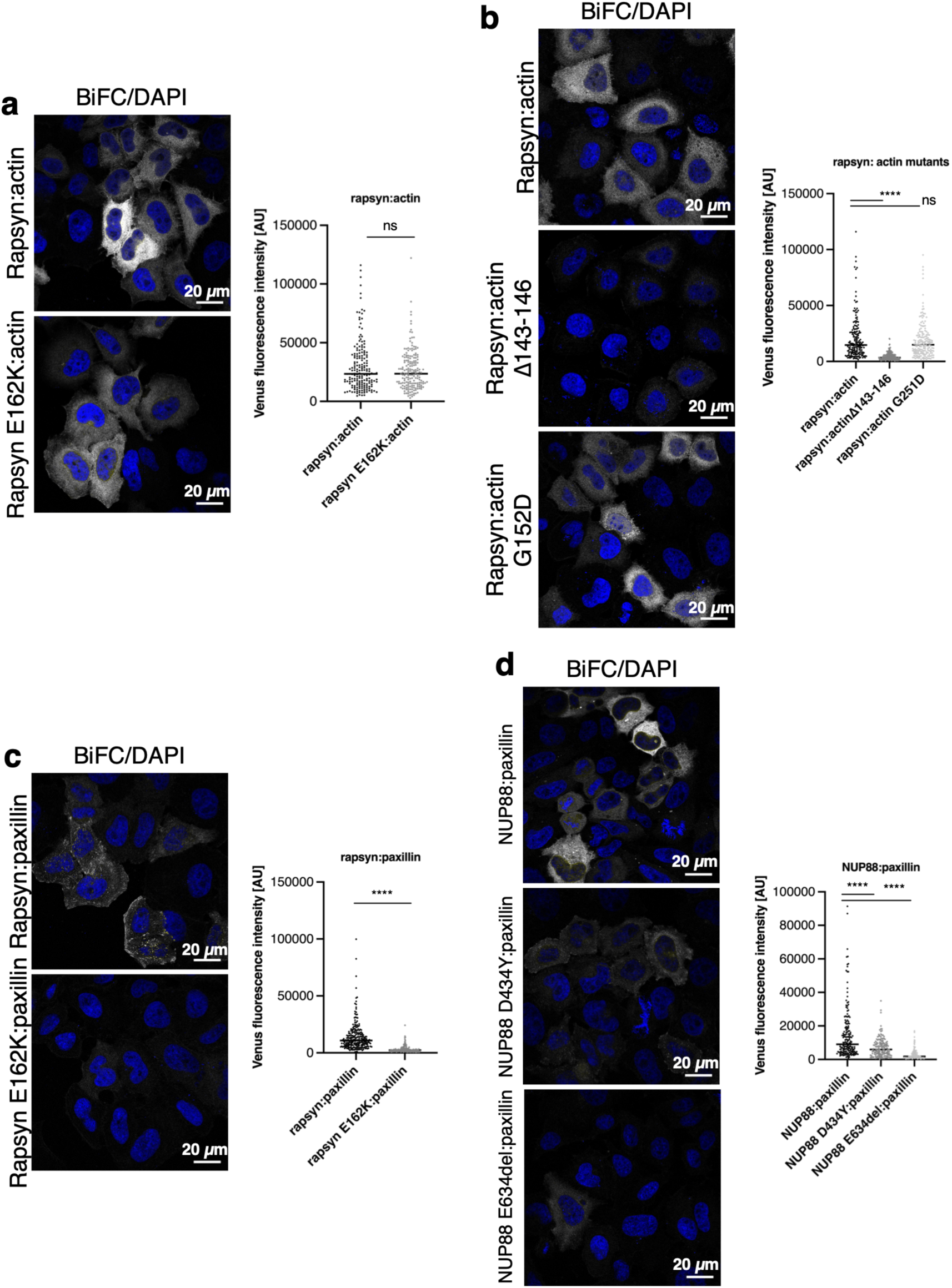
FADS-related mutations in rapsyn and NUP88 abolish their respective interaction with paxillin. Confocal microscopy analysis of BiFC signal produced by interaction between (**a** and **b**) rapsyn and actin, (**c**) rapsyn and paxillin, and (**d**) NUP88 and paxillin in HeLa cells. The FADS-related E162K mutation in rapsyn and the D434Y and E634del mutations in NUP88 largely abolished the interaction of both rapsyn and NUP88 with paxillin. Wild-type and mutant rapsyn further interact with actin and this interaction is disrupted by a mutation in actin’s hydrophobic binding cleft (actinΔ143-146), but not by a mutation affecting the polymerisation ability of actin (actinG251D). DNA was stained with DAPI (blue). Shown are representative images from at least three independent experiments. 100-200 transfected cells for each condition were analysed. Black horizontal line represents the median. *\*\*\*\*p* < 0.0001; ns, *p* not significant. Two-Way Anova test was used to calculate statistics.

To probe the biochemical interaction of rapsyn and NUP88 with FAs, we next expressed rapsyn and NUP88, respectively, paired with vinculin, but which did not result in any fluorescence signal (Supplementary Fig. S4b online). In contrast, pairs of rapsyn and NUP88 with the FA protein paxillin, produced cytoplasmic fluorescent signals (Fig. 6c,d). Rapsyn:paxillin showed the typical FA staining pattern (Fig. 6c), whereas NUP88:paxillin was more homogeneously distributed throughout the cytoplasm (Fig. 6d). FADS-related mutations in both rapsyn (Fig. 6c) and NUP88 (Fig. 6d) consistently led to a significant decrease in the fluorescence signal, suggesting that the interface of rapsyn and NUP88 with paxillin is impaired in FADS.

## DISCUSSION

The pathogenesis in fetal akinesia deformations sequence (FADS) has been described extensively and due to modern sequencing approaches several new genes have been linked to the disease^42^. In contrast to the genetic causes, a detailed molecular understanding for the pathogenesis in FADS is still lacking. Here, we use primary fibroblasts from two FADS individuals and we show that these cells have abnormal organisation in the actin cytoskeleton and increased bundling of contractile actin-myosin stress fibres (SFs). We found that the FADS proteins rapsyn and NUP88 localise nearby adhesion plaques by interacting with the focal adhesion protein paxillin and that this interaction is abolished for mutant FADS proteins.

SFs are found in many cultured non-muscle cells and are linked to the plasma membrane by focal adhesions (FAs)^43^. Formation and bundling of SFs is regulated by the small GTPase RhoA, and RhoA is also able to induce FA maturation^44^. Moreover, mature FAs can also provide a positive feedback loop in RhoA activation by serving as enrichment centre for guanine nucleotide exchange factors for RhoGTP^45, 46^. Our findings suggest that over-activation of RhoA leads to increased SF formation and FA maturation in FADS fibroblasts; in consequence, we observed that FADS cells have pronounced actin-myosin bundles and mature vinculin-positive FAs. We further propose that the alterations in the actin cytoskeleton in FADS fibroblasts lead to abnormal cellular contraction. Given that SFs in their composition resemble nascent myofibrils of skeletal muscle cells, it is tempting to speculate that the abnormal muscular contraction seen in the disease FADS arises from anomalies in the myofibril structure. Unfortunately, no post-mortem skeletal muscle biopsies from the FADS individuals investigated in this study were available for analysis of their myofibril ultrastructure. Notably, however, we previously analysed sarcomere organisation in muscle sections of a NUP88-related FADS individual^12^. Here, no changes in the gross sarcomere organisation were detected. Intriguingly, it has been shown that deficiency in lamin A, a known interactor of NUP88^47^, promotes actin-myosin filament formation and induces actin-myosin network anomalies^48^. This report supports our hypothesis that a deficiency of FADS-related proteins not only dictates actin SF anomalies but also disorganisation in the myofibril structure. Therefore, a detailed analysis of the myofibril ultrastructure in FADS is needed in future investigations.

Using proximity ligation assays (PLAs) in control foetal and in FADS fibroblasts we initially mapped rapsyn and NUP88 in proximity to vinculin. Using bimolecular fluorescence complementation (BiFC) in HeLa cells, however, revealed that both rapsyn and NUP88 are not directly interacting with vinculin, but with paxillin. This discrepancy likely arises from the different levels of resolution of the two methods: PLAs permit the detection of protein-protein associations within 40 nm distance^49^, whereas BiFC necessitates a distance from at least 7 nm^50^. Our findings confirm recent proximity-dependent biotinylation studies which identified the association of NUP88 with vinculin and paxillin^51^. Paxillin is among the first proteins recruited to newly assembled FAs^52^ and it has the largest number of potential protein binding partners within FAs^53^. Phosphorylation of paxillin by the focal adhesion kinase is a key step during FA formation. This is because it creates binding sites that serve as main binding platforms for paxillin interactors^54^ and which further promote downstream regulation of actin by FAs. Introducing FADS-related mutations into rapsyn (E162K) and NUP88 (D434Y and E634del) abolished the interaction of either with paxillin. Similar results were obtained in PLAs of vinculin and rapsyn in FADS cells carrying a homozygous E162K mutation in rapsyn. Our results suggest that the loss of the interaction of rapsyn and NUP88 with paxillin (i) leads to a disruption of a regulatory cascade at FAs and (ii) gives rise to the actin cytoskeletal anomalies seen in FADS fibroblasts. Actin-myosin contraction regulates focal adhesion maturation and recruitment of core focal adhesion proteins^55^. Therefore, future work shall address how FADS-related proteins regulate actin dynamics at FAs and which key regulatory mechanisms are involved.

Additionally, using BiFC we show that rapsyn interacts with actin possibly through the hydrophobic binding pocket of actin. FADS-related mutation E162K in rapsyn had no influence on the interaction, which can be explained by the fact that rapsyn interacts with actin-binding proteins^38, 39^ and that these interfaces might rather be affected by the mutation in rapsyn. We therefore hypothesize that a secondary mechanism for rapsyn is in place regulating actin-myosin contraction. This mechanism might further be influenced by the interaction between rapsyn and NUP88, which is abolished in FADS 2 fibroblasts^13^. However, FADS-related mutations in NUP88 appear to have no influence on the interaction with rapsyn (Supplementary Fig. S4d online), indicating that a complex interplay of the rapsyn and NUP88 with the action cytoskeleton.

Conclusively, our work establishes actin cytoskeletal anomalies as a key mechanistic factor in FADS fibroblasts possibly caused by perturbed signalling at adhesion plaques. Further investigations how these findings translate into skeletal muscle tissue organisation are needed.

## Methods

All experiments were carried out at room temperature unless otherwise stated. Analyses were performed in duplicate and were all repeated at least three times.

### Cell culture and transfection

MRC5 (AG05965-G) and FADS (FADS 1; GM11328) fibroblasts were obtained from Coriell (Human Genetic Cell Repository; Coriell Institute, Hamden, NJ, USA). FADS 2 fibroblasts were from an individual harbouring a homozygous c.484G>A (p.Glu162Lys) variant in the *RAPSN* gene (NM_005055.5) as described previously^4^. MRC5 and FADS 2 fibroblasts were grown in Minimum Essential Medium (MEM) medium (Life Technologies Gibco, Gent, Belgium) supplemented with 10% foetal bovine serum (FBS) and 1% penicillin/streptomycin (pen/strep). FADS 1 cells were grown in MEM Alpha (Lonza, Basel, Switzerland) supplemented with 15% FBS and 1% pen/strep. HeLa cells were grown in Dulbecco’s modified Eagle medium (DMEM) supplemented with 10% FBS and 1% pen/strep. All cell lines were grown at 37°C in a humidified incubator with 5% CO_2_ atmosphere.

Plasmids were transfected using TurboFect transfection reagent (ThermoFisher Scientific, Basel, Switzerland) and siRNAs using Lipofectamine RNAiMax (ThermoFisher Scientific) following the instructions of the manufacturers. siRNAs were from Dharmacon (Lafayette, CO, USA): *RAPSN* (L-006550-00), *NUP88* (L-017547-01-0005), and non-targeting siRNAs (D-001810-10).

### Plasmids

Plasmids pDEST-ORF-V1 and pDEST-ORF-V2 (Addgene plasmids #73637 and #73638) were a gift of Darren Saunders (The Kinghorn Cancer Institute, Sydney, Australia). Rapsyn-V1 and rapsyn-V2 were generated by Gateway cloning using pDONR223-RAPSN (ABIN5316678; Genomics online; GenBank: BC004196.2), as described in^13^. All other constructs were generated by In-Fusion Cloning (Takara, Saint-Germain-en-Laye, France) using the manufacture’s instructions. Mutations were inserted by site-directed mutagenesis using the QuikChange Lightning site-directed mutagenesis kit (Agilent Technologies, CA, USA) following the manufacturer’s instructions, deletions were inserted by In-fusion cloning. All constructs were verified by DNA sequencing. Plasmids used in this study are listed in Supplementary Table S1, primers are listed in Supplementary Table S2.

### Immunofluorescence

Cells were grown on crossbow-shaped micropatterns (size M; CYTOO Inc., Grenoble, France) or glass coverslips in 6-or 24-well plates, respectively, and fixed with 4% PFA in PBS for 5 min, permeabilised with 0.5% Triton-X 100 in PBS for 5 min and then fixed again. For cell spreading experiments, fibroblasts were seeded on coverslips in 24-well plates and fixed after 3 h. Blocking was performed with 2% BSA/0.1% Triton-X 100 in PBS for 30 min. Cells were incubated with the primary antibody over-night at 4°C in a humidified chamber, washed three times with 0.1% Triton-X 100 in PBS for 5 min and incubated with the secondary antibodies for 1 h in a humidified chamber, washed again and mounted with a drop of Mowiol-4088 (Sigma-Aldrich, St. Louis, MO, USA) containing DAPI (1 µg/ml; Sigma-Aldrich). Cells were imaged using a Zeiss AxioObserver.Z1 microscope (Zeiss, Oberkochen, Germany) or a Leica TCS SP8 confocal laser scanning microscope (Leica Microsystems, Heerbrugg, Switzerland). Images were recorded using the microscope system software and processed using Fiji/ImageJ, version 1.54c^56^.

The following antibodies were used as primary antibodies: mouse anti-vinculin (1:400; clone hVIN-1, Sigma), rabbit anti-non-muscle myosin heavy chain II-A (NMIIA) (1:1000; PRB-440P-050, Eurogentec, Liege, Belgium), mouse anti-α-actinin (1:800; clone EA53, Sigma), rabbit antimyosin light chain (phospho S20; MLC-P; 1:400; ab2480, Abcam, Cambridge, UK) and rabbit anti-rapsyn (1:200; NBP1-85537, Novus Biologicals, Abingdon, UK). Phalloidin-Alexa Fluor 488 (1:1000; A12379, Life Technologies Invitrogen) and phalloidin-iFluor 594 (1:1000; ab176757, Abcam) were used to stain F-actin. Secondary antibodies were goat anti-mouse IgG Alexa Fluor 488 and Alexa Fluor 568 as well as goat anti-rabbit IgG Alexa Fluor 488 and Alexa Fluor 568 (1:1000; Life Technologies Invitrogen).

### Micropattern analyses

A schematic overview of the micropattern analysis procedure is illustrated in Supplementary Fig. S1 online. First, immunofluorescence images of fibroblasts grown on crossbow-shaped micropatterns were automatically analysed using a macro re-written in ImageJ (based on a macro developed by Centre Commun de Quantimétrie (Lyon, France) and CYTOO Inc.). Briefly, images were filtered (Gaussian blur), threshold was set (Median, Huan dark), and micropatterns were centred. Images with multiple nuclei and mitotic cells were automatically excluded, stacks in each colour channel were aligned using plugins MultiStackReg (version 1.45) and TurboReg (version 2.0.0). Finally, a reference cell for each colour channel using Z-projection with a Rainbow RGB colour-coded Lookup Table was created (Supplementary Fig. S1, 1-7 online).

Next, actin filament organisation (branches and junctions) as well as actin and focal adhesion orientation was investigated in ImageJ using plugins Analyze Skeleton (2D/3D; version 3.3.0) and OrientationJ Analysis (version 2.0.4), respectively. Phalloidin and vinculin immunofluorescence images of fibroblasts after filtering and alignment of the stacks were used as input files (Supplementary Fig. S1, 8-9 online). Further, in order to examine actin filament and focal adhesion properties in more detail, we used the open-source JRE analysis tool FilamentSensor (version 0.1.7)^30^. FilamentSensor detects length, width and location (centre X and Y coordinates in an image-specific coordinate system: the x-axis is the horizontal border and the y-axis is the vertical border of the input image) for each single filament of an image. Phalloidin and vinculin immunofluorescence images of fibroblasts after filtering and alignment of the stacks were used as input files (Supplementary Fig. S1, 10 online). NMIIA organisation was analysed as described^57^. Shortly, in ImageJ a 150 px line scan was drawn across NMIIA immunofluorescence staining in actin arcs (and ventral stress fibres; data not shown) and the fluorescence plot profile was obtained. The NMIIA peak frequency was determined in R (version 3.6.0; RStudio, version 1.2.1335) using package ggpmisc (version 0.3.1) and splus4R (version 1.2-2). NMIIA immunofluorescence images of fibroblasts after filtering and alignment of the stacks were used as input files (Supplementary Fig. S1, 11 online).

### Rho GTPase assay

Active Rho Detection kit (#8820; Cell Signaling Technology/Bioke, Leiden, The Netherlands) was used to determine activation of Rho GTPase in the fibroblasts. Cells were harvested under non-denaturing conditions as described by the manufacturer, and 500 μg of lysate was either subjected to treatment with 0.1 mM GTPγS (positive control) or directly processed for affinity precipitation of GTP-bound Rho. Lysate, glutathione resin and GST-Rhotekin-RBD were incubated for 1 h at 4°C with gentle rocking. Samples were eluted with 2x reducing sample buffer containing 200 mM dithiothreitol, and incubated for 5 min at 95°C. The eluates were subsequently loaded on 12% polyacrylamide gels and Western blotting was carried out.

### Western blot

Cells were lysed in lysis buffer (10 mM Tris-HCl, pH 7.5, 150 mM NaCl, 0.5 mM EDTA, 0.5% Nonidet-P40, protease-phosphatase inhibitor cocktail tablets) and 30 µg of protein were loaded and separated by sodium dodecyl sulphate-polyacrylamide gel electrophoresis (SDS-PAGE). The proteins were transferred onto a PVDF membrane (Immobilon-P, Merck Millipore, Massachusetts, USA) and the membranes were blocked with PBS containing 5% non-fat dry milk for 30 min. The membranes were then incubated over-night at 4°C in blocking solution containing a primary antibody followed by washing with TBS containing 0.1% Tween 20 for 30 min. The membranes were next incubated with secondary antibodies for 1 h, washed 1 h in TBS containing 0.1% Tween 20 and developed. X-ray films were scanned and processed using ImageJ.

The following antibodies were used as primary antibodies: rabbit anti-RhoA (1:667; #8789, Cell Signaling Technology) and rabbit anti-GAPDH ((14C10) 1:1000; #2118, Cell Signaling Technology). Secondary antibodies were alkaline phosphatase-coupled IgG antibodies (1:10000; Sigma-Aldrich).

### Proximity ligation assay

Fibroblasts grown on cover slips were fixed with 4% PFA in PBS for 5 min, permeabilised with 0.5% Triton X-100 in PBS for 5 min and then fixed again. Blocking was performed with 2% BSA/0.1% TritonX-100 in PBS for 30 min.

All antibodies used for proximity ligation assay (PLA) were diluted in blocking solution. Antibodies mouse anti-vimentin, rabbit anti-vimentin (1:100), mouse anti-vinculin, mouse anti-NUP88, rabbit anti-NUP88 and rabbit anti-rapsyn were incubated at 4°C over-night in a humidified chamber. Phalloidin-Alexa Fluor 488 was used to visualise F-actin. Excess antibodies were removed by three washing steps using 0.1% Triton X-100 in PBS for 5 min. PLA was performed using Duolink® PLA Fluorescent Detection (red) with anti-mouse PLUS and anti-rabbit MINUS oligonucleotides (Sigma-Aldrich). PLA was performed as described elsewhere (Lin et al., 2015).

Wash buffers were prepared as follows: wash buffer A (0.01 M Tris, pH 7.4, 0.15 M NaCl, 0.05% Tween 20) and wash buffer B (0.2 M Tris, pH 7.5, 0.1 M NaCl). Cover slips were mounted onto microscope slides with Mowiol-4088 containing DAPI. Cells were imaged using a Zeiss AxioObserver.Z1 microscope. Images were recorded using the microscope system software and processed using ImageJ. PLA foci were counted using a macro written in ImageJ: briefly, parameters, scale and image threshold were set (Median, Gaussian blur, MaxEntropy) and particles (PLA foci) counted.

### Bimolecular fluorescence complementation assay

HeLa cells were grown in 24-well plates on cover slips and transfected with 200 ng of each plasmid using Turbofect transfection reagent. After 24 hours, cells were fixed in 2% formaldehyde in PBS for 15 min, washed in PBS, permeabilised with 0.2% Triton X-100 in PBS containing 2% bovine serum albumin for 10 min. Blocking was performed with 2% BSA in PBS for 30 min and cells were incubated with phalloidin-iFluor 594 reagent (1:1000; ab176757; Abcam) for 1 hour.

After three washes in PBS, samples were mounted on glass slides using Mowiol containing 1 µg/ml DAPI and stored at 4°C until viewed. Cells were viewed using a confocal laser scanning microscope (Leica TCS SP8, Leica Microsystems, Heerbrugg, Switzerland). Images were recorded using the microscope system software with uniform laser settings for all samples. Fluorescence intensity was measured using the Fiji/Image J. Images were processed using Fiji/Image J and Adobe Photoshop.

### Statistics

Plots and statistics were created and calculated using R (version 4.2.3, RStudio 2023.03.0 Build 386) using package ggplot2 (version 3.4.1) and ggstatsplot (version 0.11.0).

Dunn’s non-parametric all-pairs comparison test for Kruskal-type ranked data has been performed to calculate statistics. Holm-Bonferroni method has been used to adjust *p-*values for multiple comparisons. For all statistical tests reported in the plots the following characteristics are given: significance (*p*_Holm-adj;_ confidence interval 99%), median (µ_median_) and number of observations (n).

Alternatively, plots and statistics were established using GraphPad Prism 9 (GraphPad Software, Boston, MA, USA). Two-Way Anova tests were carried out to determine *p*-values.

### Image design

Figures were created using Adobe Photoshop (version 12.0), Omnigraffle (Omni Group, Seattle, WA, USA), and Inkscape 1.2. Colour panels were adapted from colorbrewer2.org and from the R package viridis (version 0.6.2). Colour-blind correction of RGB images was done in ImageJ using plugin Colorblind Actionbar.

## Acknowledgements

The authors thank Dr. Darren Saunders (The Kinghorn Cancer Institute, Sydney, Australia) for sharing reagents and Antonia van den Broek for technical assistance. The rabbit NUP88-antibody was a kind gift from Dr. Ulrike Kutay (ETH Zurich, Switzerland). Confocal images were acquired at the Imaging Core Facility, Biozentrum, University of Basel. This work was supported by a research grant to BF from the Fonds de la Recherche Scientifique-FNRS Belgium (grant numbers J.0102.18), as well as by the Fédération Wallonie-Bruxelles (ARC 4.110. F.000092F).

## Author contributions

Conceptualisation, R.J. and B.F.; methodology, R.J., V.M., C.R., B.F.; software, formal analyses and writing—original draft preparation, R.J.; writing—review and editing, R.J. and B.F.; supervision, B.F.; project administration, B.F.; funding acquisition, B.F. All authors have read and agreed to the published version of the manuscript.

## Data availability statement

All data generated or analysed during this work are included in this published article.

## Additional Information

The authors declare no competing interests.

