## Supplementary figures and tables for "Alteration of actin cytoskeletal organisation in fetal akinesia deformation sequence"

#### Supplementary information

##### Supplementary figures

**Figure S1. Workflow of micropattern analysis.** Each analysis step is numbered. (**1-7**) Immunofluorescence image stacks were automatically analysed using a macro written in ImageJ. Images were filtered, threshold was set and images centred. Images were filtered for single nuclei and interphase cells. Stacks were aligned using plugins MultiStackReg and TurboReg. Finally, a reference cell for each colour channel was created using Z-projections with a Rainbow RGB colour-coded Lookup Table. (**8-9**) Actin filament organisation (branches and junctions) as well as actin and focal adhesion orientation was investigated in ImageJ using plugins Analyze Skeleton and OrientationJ Analysis, respectively. (**10**) Actin filament and focal adhesion properties were analysed in detail by FilamentSensor[1]. (**11**) NMIIA organisation was analysed as described[2].

**Figure S2. Non-muscle myosin peak frequency and the number of actin branches and junctions are unchanged in FADS fibroblasts.** Violin plots of (**a**) NMIIA peak frequencies in actin arcs, (**b**) NMIIA peak frequencies in ventral stress fibres, (**c**) actin branches per cell, and (**d**) actin junctions per cell. NMIIA peak frequencies were analysed as described before[2]. Actin branches and junctions were measured in ImageJ using plugins Analyze Skeleton. n denotes the number of observations and μ the median (red dot). px, pixel.

**Figure S3. Negative controls for proximity ligation assays.** PLAs were either conducted without primary antibodies (probes only) or with each primary antibody only (anti-rapsyn, anti-vinculin, anti-NUP88).

**Figure S4. Bimolecular fluorescence complementation assays.** Confocal microscopy analysis of BiFC signal produced by between (**a**) NUP88:actin and rapsyn:actin, (**b**) NUP88:vinculin and rapsyn:vinculin, and (**c**) NUP88:rapsyn in HeLa cells. The deletion Δ346-349 in actin modifies its hydrophobic binding cleft and the E167A and S350A mutations substitute amino acids surrounding the binding cleft. The D434Y and E634del mutations in NUP88 are FADS-related. DNA was stained with DAPI (blue). Shown are representative immunofluorescence images from at least three independent experiments. (**c**) Approximately 150 transfected cells were analysed per condition. ns, *p* not significant. Two-Way Anova test was used to calculate statistics.

Figure S1

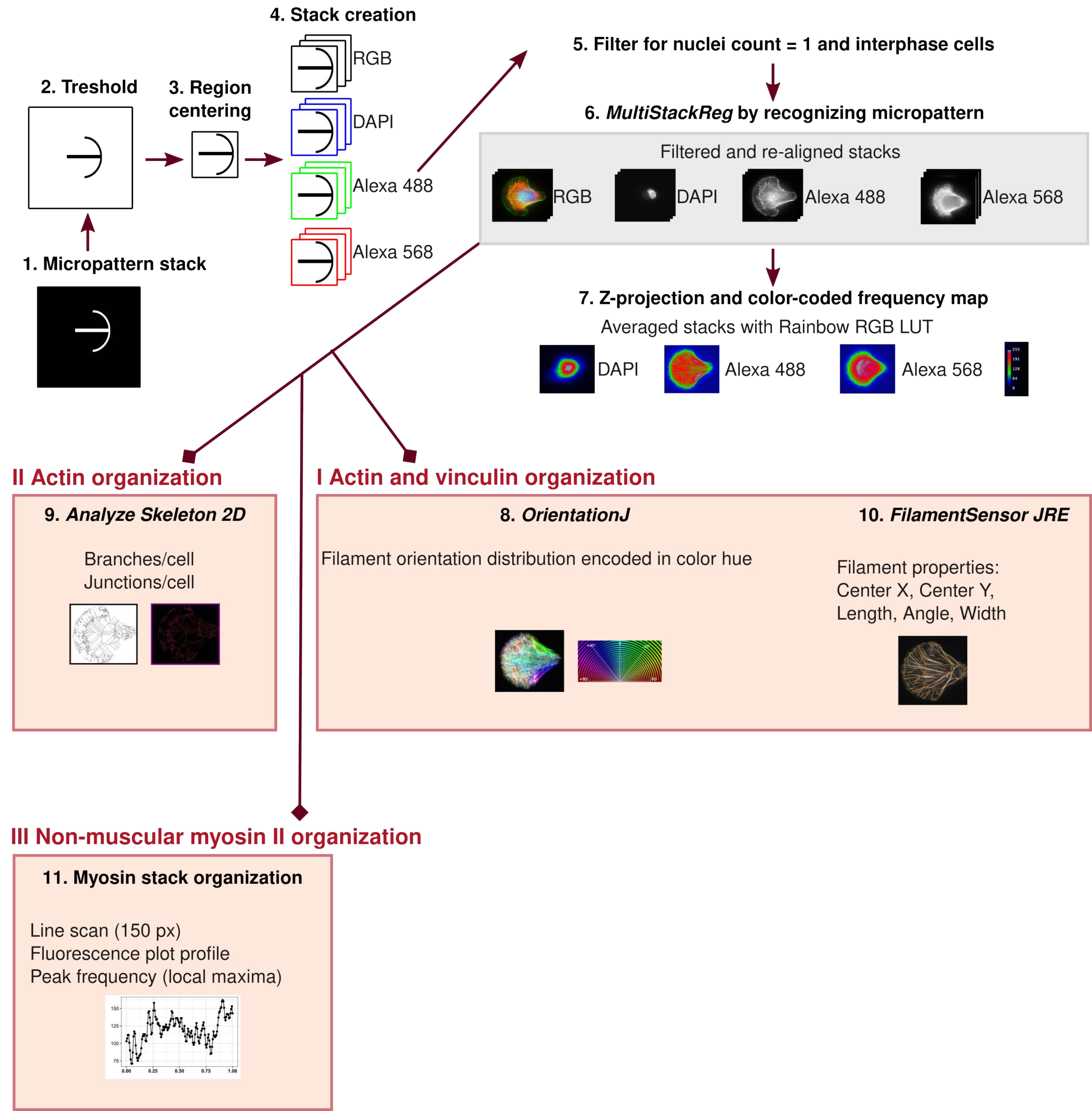

1. Micropattern stack
2. Treshold
3. Region centering
4. Stack creation
RGB
DAPI
Alexa 488
Alexa 568
5. Filter for nuclei count = 1 and interphase cells
6. MultiStackReg by recognizing micropattern
Filtered and re-aligned stacks
RGB
DAPI
Alexa 488
Alexa 568
7. Z-projection and color-coded frequency map
Averaged stacks with Rainbow RGB LUT
DAPI
Alexa 488
Alexa 568
II Actin organization
9. Analyze Skeleton 2D
Branches/cell
Junctions/cell
I Actin and vinculin organization
8. OrientationJ
Filament orientation distribution encoded in color hue
10. FilamentSensor JRE
Filament properties:
Center X, Center Y,
Length, Angle, Width
III Non-muscular myosin II organization
11. Myosin stack organization
Line scan (150 px)
Fluorescence plot profile
Peak frequency (local maxima)

### Figure S2

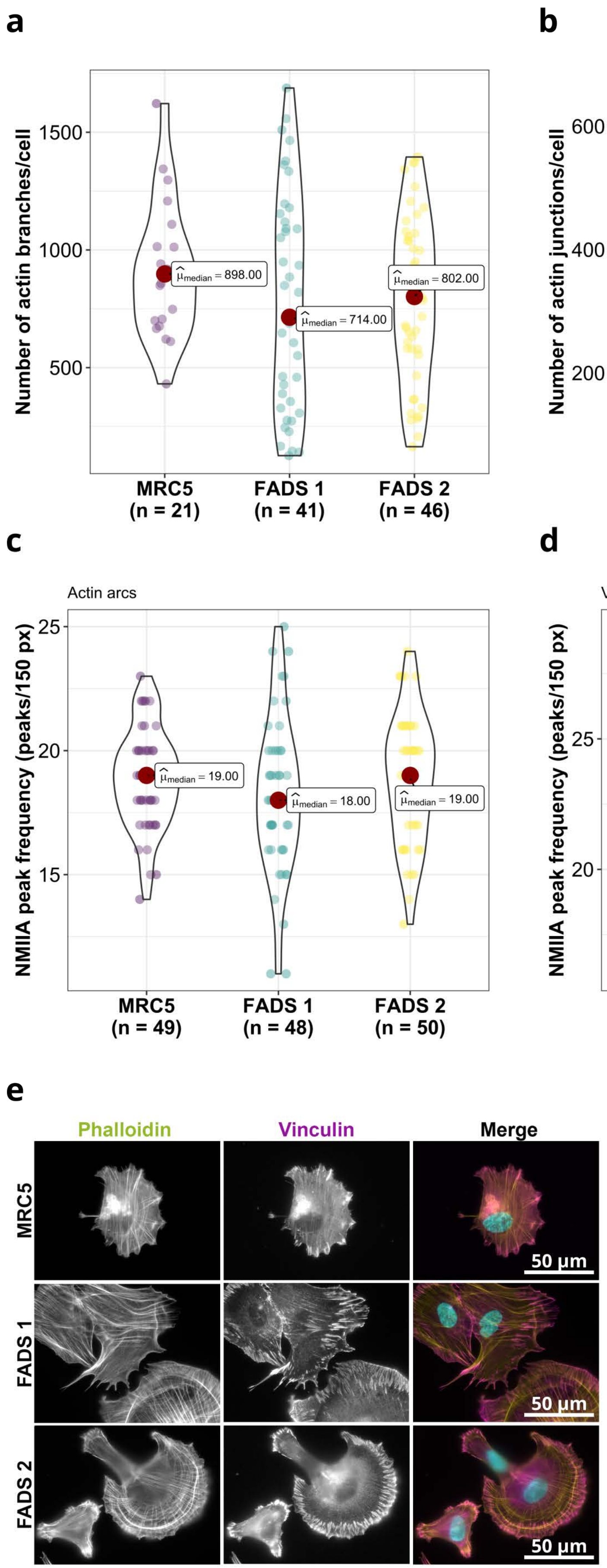

a
Number of actin branches/cell
1500
1000
500
$\hat{\mu}_{median} = 898.00$
$\hat{\mu}_{median} = 714.00$
$\hat{\mu}_{median} = 802.00$
MRC5 (n = 21)
FADS 1 (n = 41)
FADS 2 (n = 46)
b
Number of actin junctions/cell
600
400
200
$\hat{\mu}_{median} = 388.00$
$\hat{\mu}_{median} = 332.00$
$\hat{\mu}_{median} = 319.50$
MRC5 (n = 21)
FADS 1 (n = 41)
FADS 2 (n = 46)
c
Actin arcs
NMIIA peak frequency (peaks/150 px)
25
20
15
$\hat{\mu}_{median} = 19.00$
$\hat{\mu}_{median} = 18.00$
$\hat{\mu}_{median} = 19.00$
MRC5 (n = 49)
FADS 1 (n = 48)
FADS 2 (n = 50)
d
Ventral stress fibers
$p_{Holm-adj.} = 0.02$
NMIIA peak frequency (peaks/150 px)
25
20
$\hat{\mu}_{median} = 22.00$
$\hat{\mu}_{median} = 21.00$
$\hat{\mu}_{median} = 22.00$
MRC5 (n = 45)
FADS 1 (n = 47)
FADS 2 (n = 50)
e
Phalloidin
Vinculin
Merge
MRC5
FADS 1
FADS 2
50 µm
50 µm
50 µm

### Figure S3

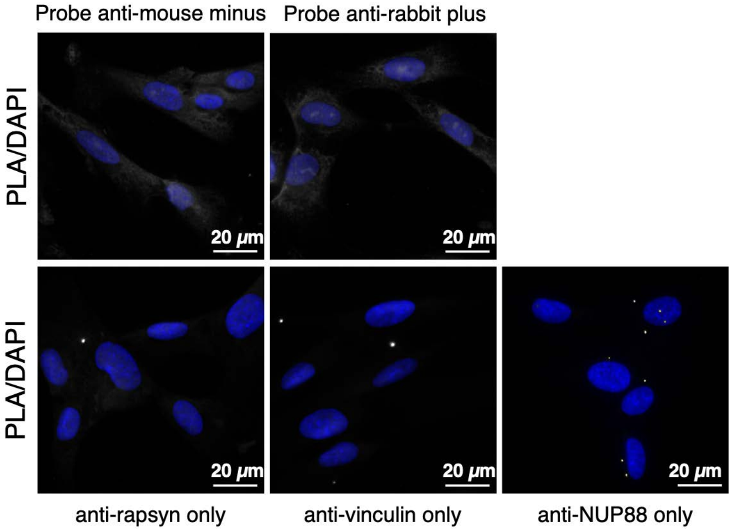
Probe anti-mouse minus
Probe anti-rabbit plus
PLA/DAPI
20 μm
20 μm
PLA/DAPI
20 μm
20 μm
20 μm
anti-rapsyn only
anti-vinculin only
anti-NUP88 only

### Figure S4

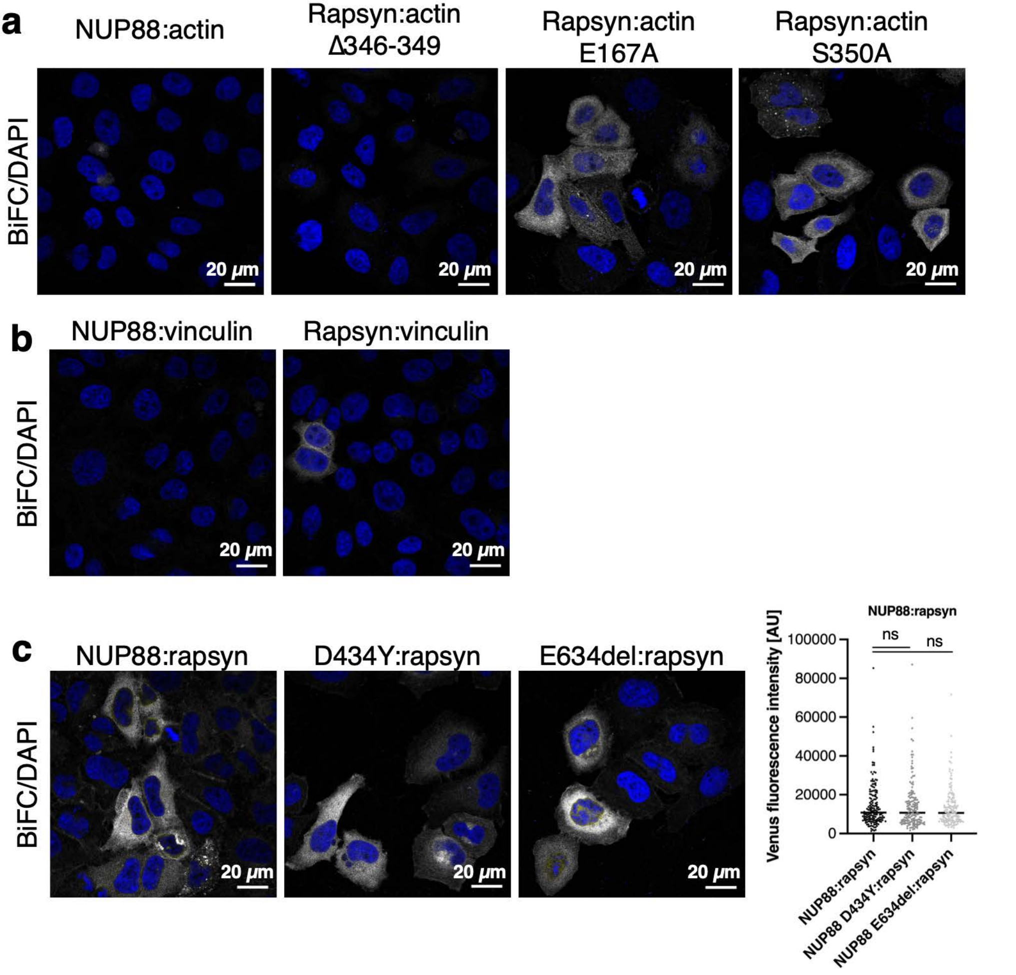
a
NUP88:actin
Rapsyn:actin
Δ346-349
Rapsyn:actin
E167A
Rapsyn:actin
S350A
BiFC/DAPI
20 μm
b
NUP88:vinculin
Rapsyn:vinculin
BiFC/DAPI
20 μm
c
NUP88:rapsyn
D434Y:rapsyn
E634del:rapsyn
BiFC/DAPI
20 μm
NUP88:rapsyn
Venus fluorescence intensity [AU]
100000
80000
60000
40000
20000
0
ns
ns
NUP88:rapsyn
NUP88 D434Y:rapsyn
NUP88 E634del:rapsyn

**Supplementary Table S2**: Primers used in this study

| Target | Sense | Sequence |
|---|---|---|
| pDEST-ORF-V1/V2 | FWD | AACCCAGCTTTcttgtacaaagtggtcatc |
| | REV | TAAGCCTGCTTTTTTGTACaaacttgtctc |
| NUP88-V1 | FWD | cAAAAAAGCAGGCTTAatgGCGGCCGCCGAGGgaccggtg |
| | REV | caagAAAGCTGGGTTGAAGTTTACATGATTGCGGATATCATTG |
| NUP88 D434Y-V1 | FWD | CTTGGATCAGATGAAGAAtATAAGGATAGTTTACAGGAACTC |
| | REV | GAGTTCCTGTAAACTATCCTTATaTTCTTCATCTGATCCAAG |
| NUP88 E634del-V1 | FWD | cAAAAAAGCAGGCTTAatgGCGGCCGCCGAGGgaccggtg |
| | REV | caagAAAGCTGGGTTGAAGTTTACATGATTGCGGATATCATTG |
| Rapsyn-E162K-V1 | FWD | caatgatgacgccatgctcAagtgccgcgtgtgctgcagcctg |
| | REV | caggctgcagcacacgcggcactTgagcatggcgtcatcattg |
| Actin-V2 | FWD | AAAGCAGGCTTAatggatgatgatatcg |
| | REV | caagAAAGCTGGGTTgaagcatttgcggt |
| Actin G251D-V2 | FWD | gacggccaggtcatcaccattgAcaatgagcggttccgctg |
| | REV | cagcggaaccgctcattgTcaatggtgatgacctggccgtc |
| Actin E167A-V2 | FWD | cactgtgcccatctacgcggggtatgccctcccccatg |
| | REV | catgggggagggcataccccgcgtagatgggcacagtg |
| Actin S350A-V2 | FWD | gctccatcctggcctcgctggccaccttccagcagatgtg |
| | REV | cacatctgctggaaggtggccagcgaggccaggatggagc |
| Actin Δ143-146-V2 | FWD | gtgctatccctgtacaccactggcatcgtgatggac |
| | REV | gtacagggatagcacagcctgg |
| Actin Δ346-349-V2 | FWD | ggcggctccatcctgaccttccagcagatgtggatc |
| | REV | caggatggagccgccgatccac |
| Paxillin-V2 | FWD | AAAGCAGGCTTAatggacgacctcgacgc |
| | REV | caagAAAGCTGGGTTgcagaagagcttgaggaagcag |
| Vinculin-V2 | FWD | GTACAAAAAAGCAGGCTTAatGCCAGTGTTTCATACGCG |
| | REV | caagAAAGCTGGGTTCTGGTACCAGGGAGTCT |

**Supplementary Table 1**: Plasmids used in this study

| Plasmid | Construct | Source |
|---|---|---|
| PBF992 | pDEST-ORF-V1 | Addgene #73637 |
| PBF993 | pDEST-ORF-V2 | Addgene #73638 |
| PBF1035 | pDEST-rapsyn-V1 | Jühlen et al., 2020 |
| PRL491 | pDEST-vinculin-V2 | This study |
| PRL492 | pDEST-actin-V2 | This study |
| PRL513 | pDEST-NUP88-V1 | This study |
| PRL556 | pDEST-NUP88 D434Y-V1 | This study |
| PRL577 | pDEST-NUP88 E634del-V1 | This study |
| PRL579 | pDEST-rapsyn E162K-V1 | This study |
| PRL581 | pDEST-actin G521D-V2 | This study |
| PRL635 | pDEST-paxillin-V2 | This study |
| PRL638 | pDEST-actin Δ143-146-V2 | This study |
| PRL639 | pDEST-actin Δ346-349-V2 | This study |
| PRL661 | pDEST-actin E167A-V2 | This study |
| PRL662 | pDEST-actin S350A-V2 | This study |
